# Low mutation load in a supergene underpinning alternative male mating strategies in ruff

**DOI:** 10.1101/2022.04.27.489720

**Authors:** Jason Hill, Erik Enbody, Huijuan Bi, Sangeet Lamichhaney, Doreen Schwochow, Shady Younis, Fredrik Widemo, Leif Andersson

## Abstract

Ruffs are shorebirds with an elaborate lekking behavior involving three male morphs with different mating strategies: Independents, Satellites, and Faeders^1,2^. The latter two are heterozygous for different versions of a supergene maintained by an inversion that were estimated to have occurred about 4 million years ago^3^. Faeders carry an intact inversion while the Satellite allele is recombinant, both of which are expected to accumulate high mutational load because they are recessive lethals. Here we have constructed a highly contiguous genome assembly of the inversion region for both the Independent and Satellite haplotypes. The recombination event(s) between an inverted and non-inverted chromosome creating the Satellite allele must have occurred recently (within the last 100,000 years) based on the minute sequence divergence between the Satellite and Independent alleles in the recombinant regions. Contrary to expectations^4,5^, we find no expansion of repeats and only a very modest mutation load on the Satellite allele in the nonrecombinant region despite high sequence divergence (1.46%). The essential centromere protein *CENPN* gene is disrupted by the inversion, and surprisingly is as well conserved on the inversion haplotypes as on the noninversion haplotype. The results suggest that the inversion may be much younger than previously thought. The lack of mutation load despite recessive lethality can be explained by the introgression of the inversion from a now extinct lineage.

---

Supergenes are defined as a cluster of functionally important polymorphisms, kept together by a structural rearrangement, usually an inversion, controlling complex phenotypic traits^4–7^. Supergenes often show a simple Mendelian inheritance and are often maintained by balancing selection. Examples of phenotypes associated with supergenes involving chromosomal inversions include Batesian mimicry in butterflies^8^, social organization in ants^6,9^, migration in trout^10^, mating systems in birds^11^, and ecological adaptation in Atlantic herring^12^ and Atlantic cod^13^. Chromosomal inversions suppress recombination by interrupting homologous recombination and present a classical paradox in genetics: how are inversions maintained over evolutionary time while interrupting fundamental processes whose disruption may negatively impact fitness? Suppression of recombination reduces the efficacy of selection to purge deleterious mutations and is expected to result in the gradual accumulation of mutational load^4,5,14^. Inversions may also accumulate repetitive elements through “degenerative expansion” that results in a larger inversion haplotype than its ancestor^4,15,16^.

Inversion polymorphisms can have low fitness in homozygotes, and in the extreme, inversions may be recessive lethals when inversion breakpoints disrupt essential genes or because deleterious mutations accumulate in a non-recombining region^4,5^. In the absence of recombination, only gene conversion and double crossovers can lead to genetic exchange between alternative chromosomal forms^17^. Thus, inversion alleles with low fitness as homozygotes are expected to harbor a high mutation load due to the absence of purifying selection in homozygotes for the lethal allele.

An old supergene inversion polymorphism is responsible for variation in male mating and plumage phenotypes of the ruff (*Calidris pugnax*)^3,18^. In this lekking species, there are three types of male mating phenotypes: Independents (Fig. 1a) have highly variable ornamental feathers, high testosterone and defend territories; Satellites (Fig. 1a) are smaller, have predominately light colored ornamental feathers and are non-territorial but take part in the lek by associating with Independents when females are present; and Faeders resemble females in body size and plumage^1,2^, and are also non-territorial. The Satellite (*S*) and Faeder (*F*) haplotypes are dominant over the Independent (*I*) haplotype^19,20^, but their allele frequencies are about 5-10% and 1%, respectively^1,3,21^. Further, both *S* and *F* are homozygous lethals, most likely because the inversion disrupts an essential centromere protein gene, *CENPN*^3,18^. *I* is ancestral, *F* is a full-length 4.5-Mb inversion that occurred about 4 million years ago, and *S* is the result of recombination between an inverted and a non-inverted allele^3^. This polymorphism is a classic example of a supergene responsible for the covariation of multiple phenotypic features. Theoretical predictions and empirical data in other systems^4,5,7,14,16,22^ indicate that the inversion alleles should be burdened by a high mutational load, but this was not possible to determine using the existing, highly fragmented, genome assemblies for ruff. How this inversion has persisted over millions of years, despite being lethal in the homozygous condition, occurring at a relatively low frequency, and carrying a predicted high genetic load is uncertain. Simulation studies suggest that mutational load can be mitigated by even moderate rates of gene conversion^16^, but this has not been demonstrated in wild populations.

**Fig. 1.**
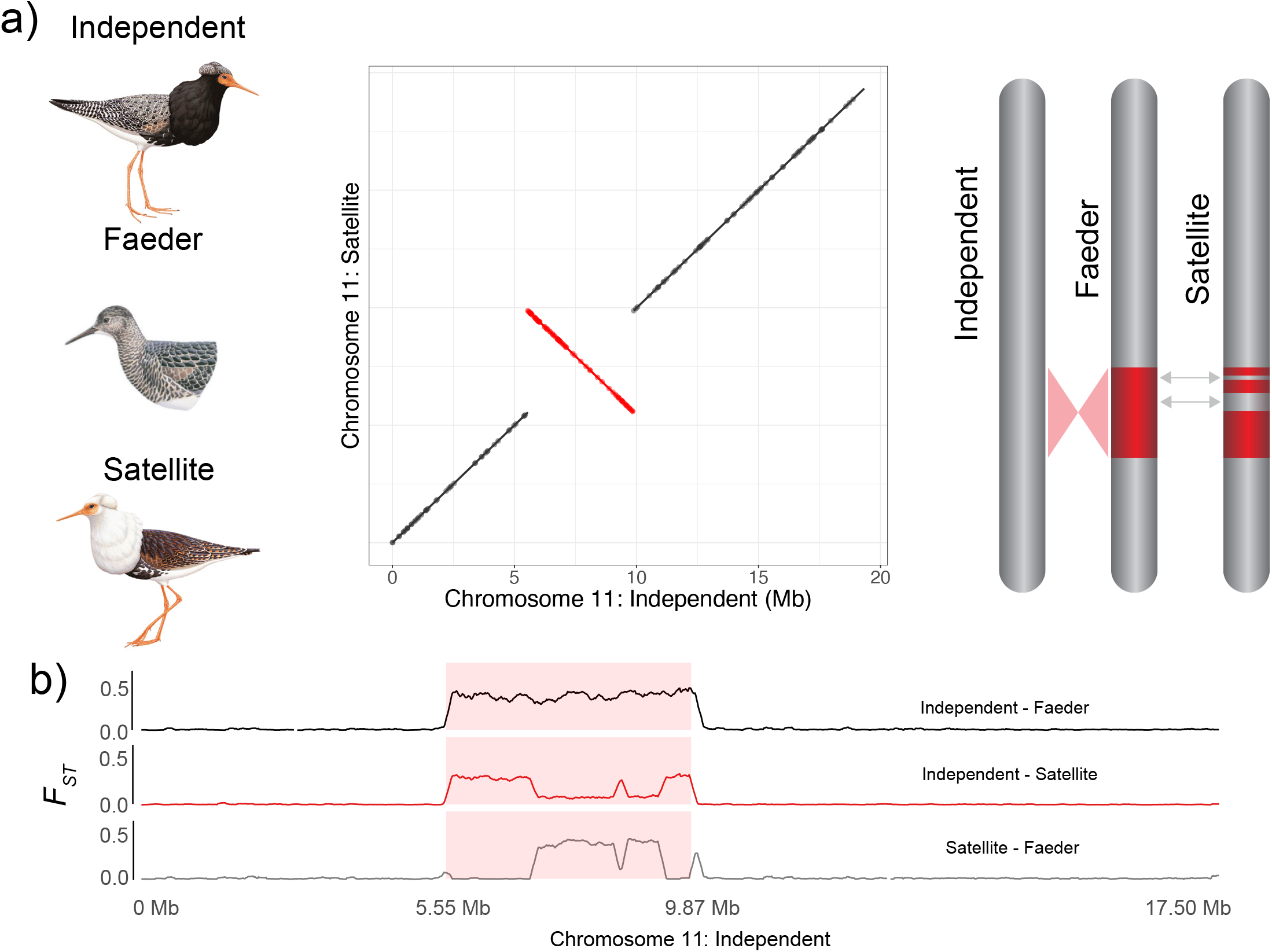
Ruff male phenotypes and inversion alleles. **a)** Left, three male ruff phenotypes: Independent, Faeder, and Satellite (illustrations reproduced with permission of Lynx edicions). Chromosome 11 alignments between Satellite and Independent assemblies with the 4.3 Mb inversion highlighted in red. Right, a graphical representation of the ancestral Independent chromosome 11, inverted Faeder chromosome, and the recombined Satellite chromosome. Satellites and Faeders are heterozygous for the inversion allele and Independents are homozygous for the non-inverted allele. **b)** Genetic divergence estimated by *F_ST_* in 15-kb rolling windows between all three chromosomal arrangements across chromosome 11. The 4.3-Mb inverted region is highlighted by a red box marking the 5.55 Mb and 9.89 Mb breakpoints.

In order to conduct detailed comparisons of molecular evolution between the inversion haplotypes, we combined a Chromium 10x linked read assembly with PacBio long read contigs to construct highly contiguous assemblies for both the Independent and Satellite haplotypes across the inverted region from one heterozygous individual. We use these assemblies and previously published whole-genome data to test the prediction that the Satellite allele shows a high mutational load and an expansion of repetitive elements. Specifically, we use estimates of synonymous and non-synonymous substitution rates to test this prediction.

## Results

### Reassembly of the region harboring the inversion polymorphism

We generated a high-quality genome assembly from a Satellite (*S/I*) individual collected in North Sweden. We first constructed a Chromium 10x linked read library which was sequenced to a depth of 90x for an estimated genome size of 1.23 Gb^3^ and conducted phase-aware genome assembly using Supernova v2.0 (Supplementary Table 1). A 17 Mb scaffold was homologous to chicken chromosome 11 and harbored the inversion polymorphism (Fig. 1a). A comparison between the two versions of this scaffold representing the two alleles identified a 4.3 Mb inversion in the Satellite allele, similar in size to the 4.5 Mb inversion as previously described^3,18^.

We sequenced PacBio long-read library from the same individual to an estimated coverage of 82x and constructed a diploid assembly using an in-house pipeline (Supplementary Fig. 1). We polished the PacBio assembly using haplotype-aware Chromium reads, which resulted in highly contiguous haplotype assemblies. We next replaced the 4.3 Mb region harboring the inversion region in the Chromium assembly with gapless genomic sequence of the matching haplotype from the PacBio assemblies (Supplementary Fig. 1). The Satellite inversion haplotype was 14 kb shorter than the corresponding Independent haplotype. Consistent with the smaller size of the Satellite haplotype, it does not include an expansion of repetitive elements (Table 1, Supplementary Fig. 2), in contrast to some other inversions where repetitive elements accumulate in the absence of recombination^4,15,22^. We identified the inversion breakpoints of the hybrid genome assembly by mapping PacBio reads and 10x reads to the final assembly (Independent: 5,548,078-9,885,008, Satellite: 5,546,737-9,869,715). We found that the Independent haplotype assembly is approximately 200 kb shorter than the previous short-read only assembly^3^. Evidence from genome and long-read alignments indicate that erroneously duplicated sequences account for ~52 kb of unique sequence in the previous assembly and 115 kb represents gap sequences. Together, these data suggest that the updated assembly reported here is both more contiguous and complete.

**Table 1.** The Satellite inversion haplotype is not enriched for repeats. Independent and Satellite chromosome 11 were aligned to each other and variants called using the software Assemblytics. In this table, each variant type refers to variants present in the Satellite haplotype called relative to the Independent chromosome. Variants are binned as in regions before, inside, or after inversion. The length of each bin is marked in Mb.

| Satellite variant | Chromosome 11 |  |  |
| --- | --- | --- | --- |
|  | Before inversion | Inside inversion | After inversion |
|  | (5.55 Mb) | (4.30 Mb) | (9.40 Mb) |
| Deletion | 15 | 30 | 16 |
| Insertion | 20 | 25 | 10 |
| Repeat contraction | 1 | 8 | 2 |
| Repeat expansion | 2 | 0 | 1 |
| Tandem contraction | 0 | 8 | 0 |
| Tandem expansion | 0 | 3 | 0 |

The gene content of the inversion of both haplotypes was annotated using transcriptome data from five previously published tissues (*n* = 10 individuals)^18^ and one new sample from skin tissue of a male Independent using the Maker (v3.01.2-beta) pipeline followed by manual refinement. We identified 107 full-length gene models, for which 100 have a known homology to annotated genes in other species, while 7 putative genes show no identifiable homology to known genes (Supplementary Table 2). These genes are supported by RNAseq data, and the ratio of nonsynonymous to synonymous substitutions (dN/dS) for four of these genes between Independent and Satellite alleles was much lower than one (mean = 0.03) indicating that those genes are under strong purifying selection. Our annotation included 13 genes that were not previously reported in the ruff inversion, one that was identified as a different ortholog, and two that were annotated previously were not found (Supplementary Table 2).

In order to evaluate the accuracy of our new assembly of the inversion region, we investigated genetic divergence and diversity using previously published^3,18^ resequencing data from 30 individuals: 16 Independents, 11 Satellites, and 3 Faeders. Genetic divergence between the Satellite, Faeder, and Independent chromosome 11 revealed sharp boundaries separating the non-recombining inversion region from the remainder of chromosome 11 in Faeder individuals (Fig. 1b). The Satellite haplotype is divided into high and low *F_ST_* regions in comparison with the Independent haplotype, which reflects the non-recombinant regions and the regions that have recombined with Independent haplotypes, respectively. The Satellite haplotype includes three distinct regions with strong genetic differentiation relative to the ancestral Faeder haplotype, but low or no genetic differentiation from the Independent haplotype (Fig. 1b). One of these regions, the smallest, is present outside of the inversion breakpoint identified by long-reads and begins at the 9.89 Mb breakpoint of the inversion and continues for 121 kb outside of the inversion. Despite being outside of the inversion, Faeders were strongly differentiated from Independents and Satellites, indicating that suppressed recombination between the Faeder and Independent chromosome extends beyond the 9.89 Mb inversion breakpoint (Fig. 2a). This region contains three genes; *NFAT5, NOB1*, and *WWP2*. This region has apparently recombined with an *I* chromosome during the formation of the *S* haplotype. A notable exception to the genetic similarity between the Independent and Satellite haplotypes in this region is a 2,082 bp LINE insertion at the 9.89 Mb inversion breakpoint in the Satellite chromosome.

**Fig. 2.**
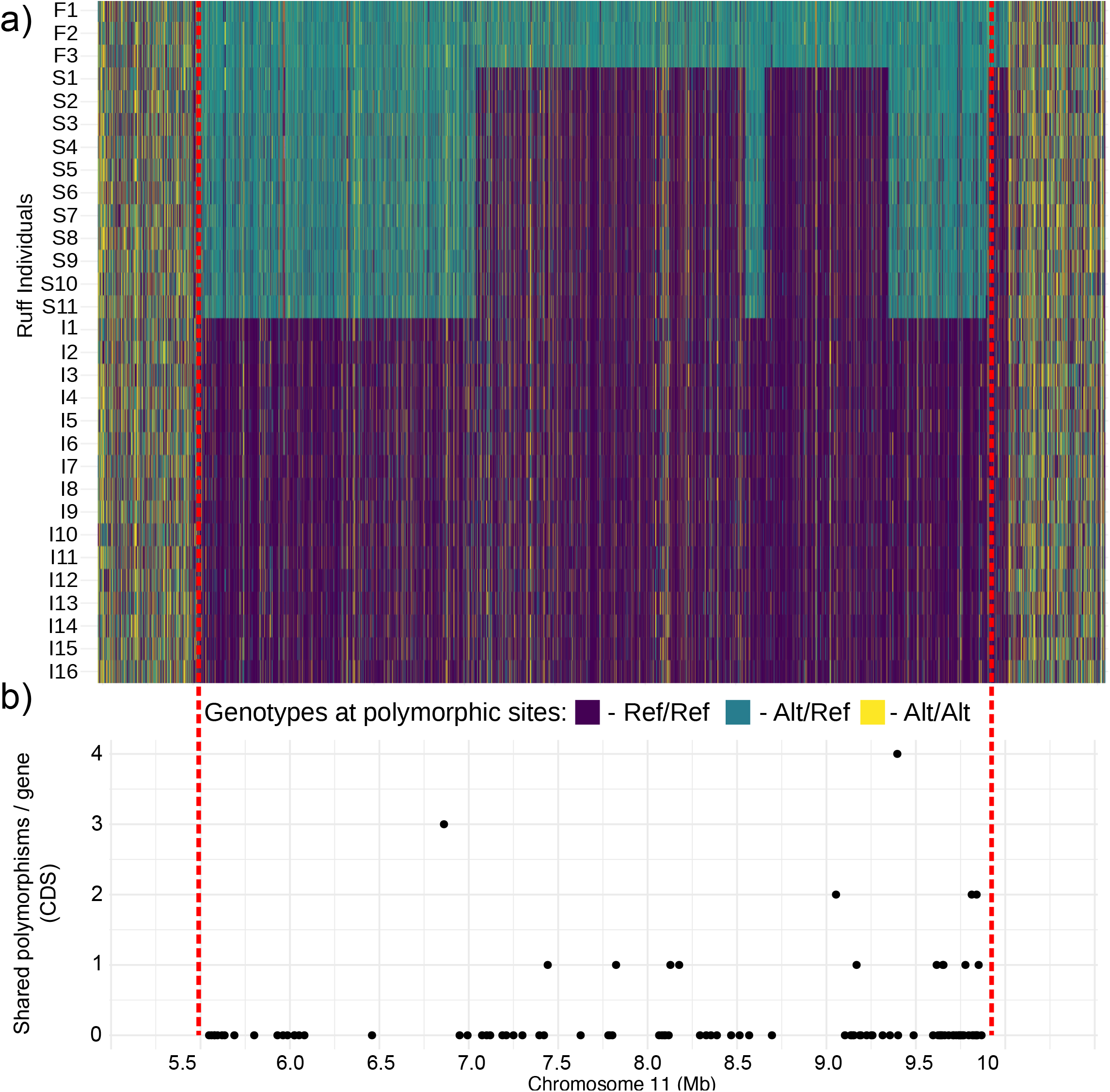
SNP genotypes in three male morphs in ruff across the inverted region. **a)** Genotypes for each SNP for the inversion region +500 kb outside each breakpoint (top panel): dark blue – homozygous for Independent reference allele, light blue – heterozygous for a reference and alternate allele, and yellow – homozygous for an alternate allele. **b)** Number of shared SNPs between Satellite and Independent chromosomes for each gene within the inverted regions. These were identified as SNPs homozygous among Satellites (*S/I*). Such shared SNPs are not expected to be present unless there is genetic exchange between the two haplotypes.

### Estimating the age of the inversion and the Satellite haplotype

We estimated the time of divergence between the Independent and Satellite haplotypes based on the net number of nucleotide substitutions (*d_a_*) between Satellite (*S*) and Independent (*I*) haplotypes. *d_a_* was calculated separately for the high and low *F_ST_* regions under the assumption that the low *F_ST_* regions are more recently diverged from each other due to recombination events occurring since the initial formation of the inversion. Within the high *F_ST_* regions, we estimated sequence divergence (*d_a_*) at 1.46% based on the average number of heterozygous sites in *S/I* heterozygotes and *I/I* homozygotes (see Methods), resulting in an estimated age of the inversion of 3.84 million years, almost identical to the previous estimate of 3.87 million years^3^. The net number of nucleotide substitutions for the low *F_ST_* region was estimated at *d_a_*=0, because pairwise nucleotide diversity (*d_xy_*=0.0028) between Satellite and Independent haplotypes was identical to the average nucleotide diversity (*d_x_*) among Independent chromosomes (Supplementary Fig. 3). Thus, the recombination event(s) in Satellite haplotypes must have occurred much more recently (<100,000 years) than the average divergence time among Independent chromosomes, which was estimated at 740,000 years before present using the estimated *d_x_* (0.0028). In other words, the recombined Satellite haplotype in these regions is as divergent from the average Independent haplotype as a randomly drawn Independent haplotype is. The age of the Satellite haplotype was previously estimated to be about 500,000 years^3^, but our analysis here shows that the Satellite haplotype arose much more recently.

### Low mutation load in the Satellite haplotype

In order to quantify mutational load and patterns of molecular selection, we calculated nonsynonymous (dN) and synonymous (dS) substitution rates for each of the genes with alignable orthologs between the two ruff haplotypes and killdeer (*Charadrius vociferus*), a related outgroup species with an annotated genome assembly (Supplementary Table 3). We aligned amino acid and cDNA sequences for Independent and Satellite haplotypes to the corresponding sequences from killdeer. We restricted our analysis to the non-recombining regions, because it is not meaningful to describe *dN/dS* in comparison to killdeer for the regions of high sequence identity between Satellite and Independent haplotypes. We found very similar dN in Independent and Satellite orthologs compared to killdeer for the great majority of genes (Fig. 3a, Supplementary Table 3). However, there was a near significant excess of Satellite orthologs with highest dN relative to killdeer; 38 genes had higher dN in Satellites compared with 27 genes showing the opposite trend (*X*^2^= 3.1, d.f.=1, *P* = 0.08), and dN was identical for 18 genes. We next calculated the expected rate of nonsynonymous substitutions on Satellite haplotypes relative to killdeer in the absence of purifying selection (i.e., dN = dS) for each Satellite ortholog since the split from the Independent allele (see Methods) and found that 73 of the 83 Satellite orthologs had accumulated fewer nonsynonymous mutations than expected (Fig. 3b), consistent with an effect of purifying selection on the evolution of protein-coding genes on the Satellite haplotype.

**Fig. 3.**
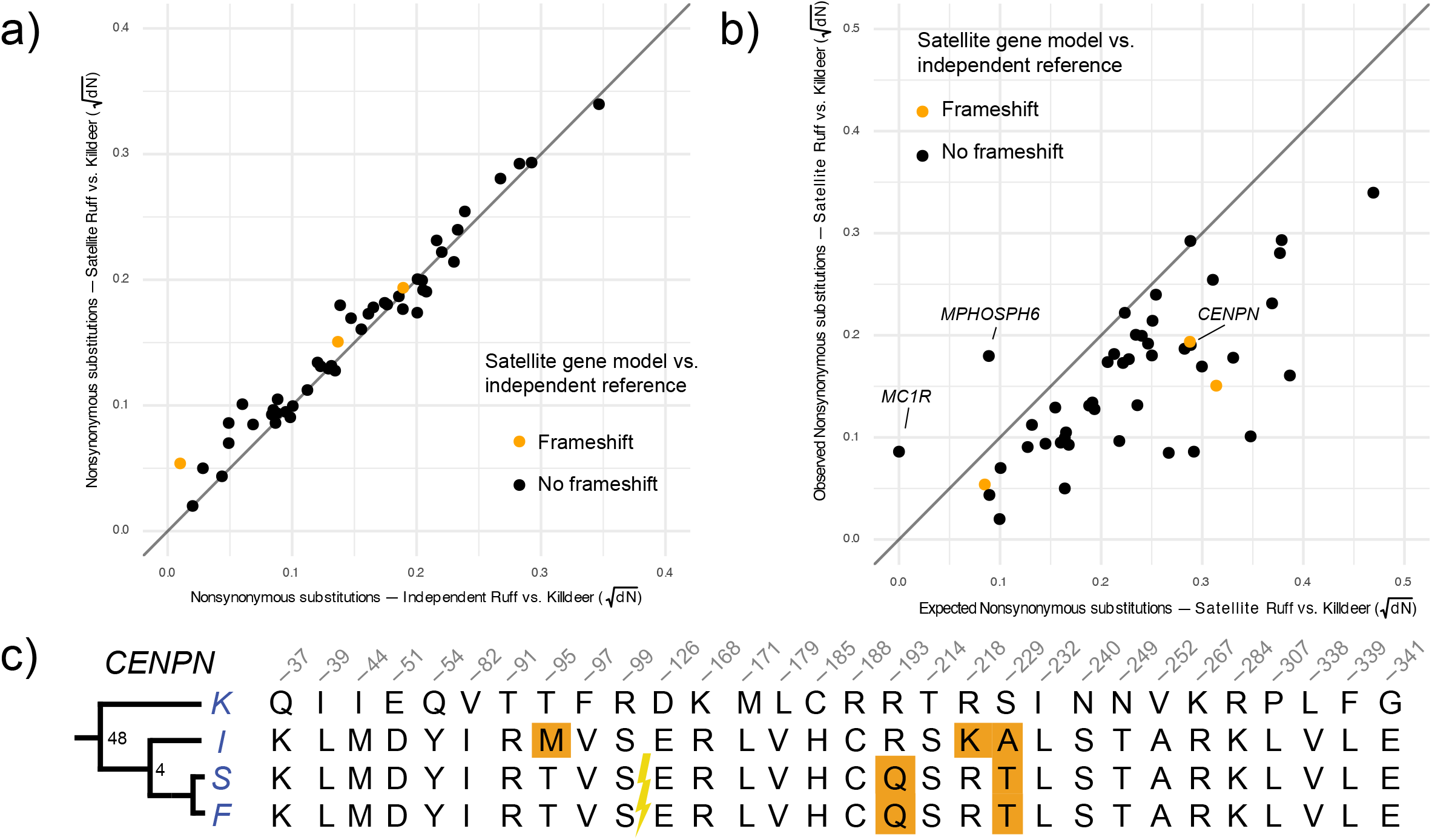
Evidence for purifying selection acting on Satellite alleles. **a)** Nonsynonymous substitution rate (dN) in pairwise comparisons of Independent or Satellite alleles *vs.* killdeer (*Charadrius vociferus)* for each gene ortholog located in regions with high *F_ST_* in the comparison of Satellites and Independents. Genes with frameshift mutations in the Satellite haplotype are marked in orange (*CMIP, TMED6*, and *COG8*), and all other genes are marked in black. **b)** The observed dN values of the Satellite *vs.* killdeer orthologs compared to expected under the assumption of no purifying selection acting on the Satellite alleles. **c**) Amino acid alignment of variable sites for the CENPN protein among killdeer and the three different ruff morphs. The exons encoding the N-terminal part of CENPN is located outside the inversion and the inversion breakpoint is indicated with a flash. The derived T95M amino acid substitution was only found among Independent chromosomes but is not a fixed difference.

The 5.55 Mb breakpoint of the inversion interrupts centromere protein N (*CENPN*) and is the predicted mechanism for the lethality of homozygous *S/S* individuals^3,18^ (Supplementary Fig. 4). We predicted that the 3’ fragment of the gene within the inversion would be a pseudogene as old as the inversion that has accumulated a roughly equal number of synonymous as nonsynonymous mutations. Unexpectedly, we found a striking reduction in nonsynonymous mutations (dN/dS = 0.18) for the inverted *CENPN* fragment and we found a very similar number of nonsynonymous substitutions (three) between killdeer and each of the three ruff haplotypes (Fig. 3c). We evaluated protein evolution under a phylogenetic context using three additional outgroups (Fig. 4a) using the Phylogenetic Analysis by Maximum Likelihood (PAML^23^) package. We found that the rate of non-synonymous substitutions in the Satellite branch did not differ significantly from the rate in the rest of the phylogeny (*P*(*ω_0_* = *ωs*) = 0.32) (Fig. 4a). This is an unlikely finding if the gene was inactivated 4 million years ago or if it has evolved a novel function subsequent to the inversion event. Furthermore, an RT-PCR-based study targeting the 3’part within the inversion using a limited set of tissues from two adult Satellite males revealed only expression of the Independent allele (Supplementary Fig. 5). Similarly, Loveland *et al*.^24^ detected only full-length transcripts from the Independent allele in Satellite heterozygotes. This does not exclude the possibility that this truncated form of CENPN is expressed in a specific tissue or during a critical stage of development.

**Fig. 4.**
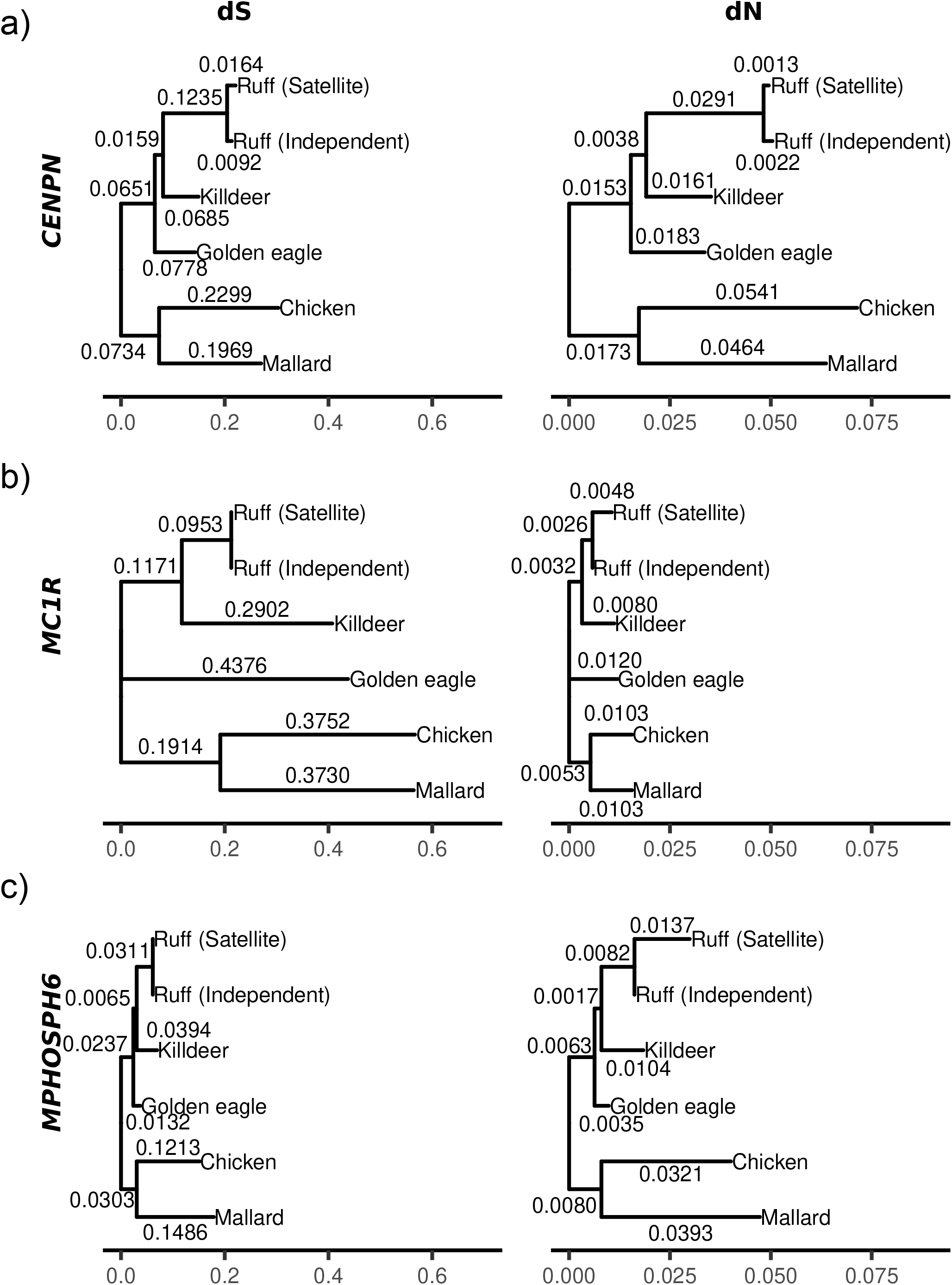
Comparison of the rate of non-synonymous substitutions in the ruff Satellite branch compared with other branches in a bird phylogeny for *CENPN (**a**), MC1R (**b**)*, and *MPHOSPH6 (**c**).* Shown here are phylogenetic trees using a model where the substitution rate ω (*dN/dS)* was allowed to vary in the ruff Satellite branch. Branch lengths correspond to dS and dN values as appropriate.

### Identification of candidate genes under selection

Two genes had a clear excess of derived amino acid substitutions on the Satellite haplotype, *melanocortin-1 receptor* (*MC1R*) and *M-phase phosphoprotein 6* (*MPHOSPH6*) Fig. 3b. Two missense mutations differ between killdeer and the ruff Independent *MC1R* alleles that share a common ancestor about 50 million years before present^25^, whereas Independent and Satellite alleles that separated about 4 million years ago differ by four (Supplementary Fig. 6). Consistent with this observation, a PAML^23^ analysis supported an accelerated protein evolution in the Satellite branch (*P*(*ω_0_* = *ωs*) < 0.001) (Fig. 4b). The observation that three of these four derived mutations in the Satellite occurred subsequent to the split from the Faeder allele (Supplementary Fig. 6) and at residues that are conserved between mammals and birds^3^ make this a clear example of how supergenes accumulate adaptive mutations subsequent to the appearance of the supergene^4,26^. *MC1R* is the obvious candidate gene underlying light colored ornamental feathers in Satellites^3^ (Fig. 1a). The Satellite *MPHOSP6* allele also carries four derived missense mutations (Supplementary Fig. 6) and not a single synonymous change (Fig. 3b; Supplementary Table 3) making this also a strong candidate gene under selection. In fact, PAML^23^ analysis supports accelerated evolution in the Satellite branch also for *MPHOSP6 (P*(*ω_0_* = *ωs*) = 0.02) (Fig. 4c). The Faeder allele shares three of these nonsynonymous changes. *MPHOSP6* is less well-studied than *MC1R*, but was identified as being phosphorylated during the M-phase in the cell cycle^27^ and is a component of the RNA exosome^28^. Genome-wide association studies in humans have found that *MPHOSP6* is associated with variation in leukocyte telomere length, a marker for genome aging^29^. Thus, the function of MPHOSP6 may be relevant for chromosome function and the evolution of the ruff supergene because the inversion disrupts the *centromere protein N* (*CENPN*) gene.

### Genetic exchange between haplotypes

Inversion haplotypes may exchange genetic material by double recombination and gene conversion^17^. Other than the three large recombinant regions described above, we did not find any evidence of frequent recombination events. In order to test the prediction of gene conversion we counted shared polymorphisms in exons within the inversion by identifying SNPs for which both homozygous genotypes were present among Satellites. This analysis identified 14 genes that carried at least one shared polymorphism (Fig. 2b) with the most frequent occurring in *Cadherin-14* (*n* = 4 shared polymorphisms). Notably, 7 of the 14 shared polymorphisms were found in the 500 kb immediately adjacent to the 9.89 Mb breakpoint. Genetic differentiation is high between Satellites and Independents in this region (Fig. 1b), suggesting that these shared polymorphisms were most likely produced through gene conversion rather than recombination.

## Discussion

Our assembly of the Satellite inversion haplotype provides an opportunity to examine the evolutionary history of chromosome segments that have been maintained in the heterozygous state, because the inversion is a recessive lethal^18^. The estimated sequence divergence of 1.46% between the non-recombining region of the Satellite haplotype and the non-inverted Independent haplotype implies that the split from a common ancestral sequence occurred about 4 million years ago. Population genetics theory predicts that the Satellite haplotype should have accumulated a high mutational load if it has been maintained as a recessive lethal over such a long time span^4,5,14^. However, we were unable to find support for this prediction.

The current understanding of the evolutionary trajectory of genomic regions that are homozygous lethal and non-recombining is towards a fate resembling that of the Y chromosome in XY sex chromosome systems^14^. In these cases, the Y chromosome accumulates deleterious mutations that inactivate most of the genes found in the homologous X chromosome. Further, the accumulation of genetic load within supergenes associated with inversions is well documented empirically in several species of plants and insects^4,8,22^. The ruff inversion polymorphism is an unexpected exception to this model. First, we see no accumulation of repetitive elements and the Satellite inversion region is about 14 kb smaller than the ancestral haplotype, due to the presence of deletions that may be adaptive^3^. Secondly, a few genes carry frameshift mutations, but the remaining 75 within the non-recombining region carry a small, if any, mutational burden in comparison to the homologous Independent chromosome. Recombination and gene conversion may cause some gene flow from Independent to variant chromosomes, but cannot explain the low dN/dS ratios in relation to the estimated time of divergence (Fig. 3b). Furthermore, given the considerable sequence divergence between haplotypes, it is surprising that the disrupted centromere protein gene *CENPN* is as well conserved on the Satellite and Faeder haplotypes as it is on the Independent haplotype containing an intact *CENPN* copy (Fig. 3c). The 3’ part of the gene that is well separated from the promotor and located inside the inversion has a much lower rate of nonsynonymous substitutions than expected if it was pseudogenized 4 million years ago or evolved an altered function subsequent to the inversion event (Fig. 3b. Fig. 4a).

The unexpected low mutational load in general and for *CENPN* in particular could be explained by efficient purifying selection for most genes present on the inversion haplotypes. Selection to maintain the protein sequence encoded by the truncated copy of *CENPN* located within the inversion assumes that is has an unknown functional role. If so, the expression of this 3’fragment of the gene must be driven by a novel promoter such as a Long Terminal Repeat (LTR). This model would gain support if expression of this fragment can be documented, but our gene expression study using adult tissues revealed expression of only the Independent allele in Satellite heterozygotes. However, an alternative and simpler explanation is that the inversion may not be as old as the sequence divergence data indicate. There are two main scenarios that would be consistent with existing data. First, that an old larger inversion has recombined much more recently, and resulted in the disruption of *CENPN* and recessive lethality, similar to the recombination events creating the Satellite haplotype (Fig. 1a) or the second inversion allele at the chicken *Rose-comb* locus^30^. Second, the inversion may be the result of an introgression event from another species. Under this model, the inversion occurred at the introgression event or after it and was favored by selection, because it kept together alleles contributing to the male mating phenotype. Such a scenario has been hypothesized for the mating-system linked supergene in white-throated sparrow (*Zonotrichia albicollis*)^31^ where introgression from an extinct species is suspected. In ruff, there are no described hybrids and the species occurs on a long branch in the Scolopaci phylogeny^32^, making an extent species an unlikely donor. We explored this possibility by searching the ruff genome for haplotypes that segregate at comparable sequence divergence as the ruff inversion and may harbor additional introgressed fragments (Supplementary Fig. 7). No other genomic regions show an elevated sequence divergence among haplotypes matching the inversion, although several small regions (<10kb) are evident and may represent regions under balancing selection. Together, the lack of a clear donor species means that it is difficult to support the introgression hypothesis, but the inversion may have originated from a now extinct lineage that hybridized with ruff in the past.

We previously estimated that the recombination event(s) responsible for the Satellite haplotype occurred about 500,000 years ago^3^. Using additional individuals and an updated genome assembly, we report a much more recent time since divergence, because nucleotide divergence between the Satellite and Independent haplotypes does not differ from nucleotide diversity among Independent haplotypes (Supplementary Fig. 3). A very similar result has been noted for inversion polymorphisms segregating in *Drosophila melanogaster*^33^. Our result implies an origin of the Satellite haplotype within the last 100,000 years. An important reason for the revised estimate is that we used a more accurate method to estimate the age of inversion haplotypes (see Methods). However, the dating of the recombination event(s) may be underestimated if genetic exchange (gene conversion and/or double recombination) between the Satellite and Independent haplotypes within the low *F_ST_* region is an ongoing process. More than one hundred missense mutations distinguish the Satellite and Independent haplotypes, but only three of them occur in regions of recombination^3^. In contrast, the great majority of missense mutations that distinguish the Satellite and Faeder haplotypes are located in the regions of recombination where Satellite and Independent haplotypes are almost identical. Thus, the Satellite haplotype constitutes a combination of one set of genetic variants shared with Independents and another set shared with Faeders, which is matched by the shared ornamental feathers between Satellites and Independents, but low levels of testosterone in blood in both Satellites and Faeders^18^.

In Satellites, males have light-colored ornamental feathers (Fig. 1a) and mutations in the *melanocortin-1 receptor* (*MC1R*) gene, located within the inversion, have been proposed to cause this phenotype^3^. Here, we found accelerated protein evolution of the *MC1R* allele in Satellites, which suggests that non-synonymous mutations in this gene have been under selection. Accelerated evolution involving multiple changes is a likely scenario because the Satellite *MC1R* allele cannot be a simple loss-of-function allele that just reduces pigmentation intensity. Firstly, loss-of-function *MC1R* alleles are recessive, such as *recessive yellow* in mouse (http://www.informatics.jax.org/marker/MGI:99456) and *chestnut* in horses^34^. Secondly, the causal allele in Satellites must have a dominant effect on ornamental feathers, but a completely recessive effect when the non-breeding feathers molt. The Satellite males are indistinguishable from Independent males outside the breeding season. A plausible scenario is that the Satellite *MC1R* allele carries a combination of regulatory mutations causing differential expression between ornamental and normal feathers, and coding changes that inhibit MC1R signaling which result in light-colored ornamental feathers.

Together we show that the evolutionary history of the ruff inversion, which determines alternate male mating strategies, is more complex than previously thought. Molecular dating suggests that inversion is very old (4 MYA), but polymorphism data suggests that it has not generated high mutation load, which would be expected for a recessive lethal supergene. A possible explanation for this apparent contradiction is that the inversion introgressed from an extinct species much more recently, or alternatively that the current organization with disruption of *CENPN* are relatively young. Furthermore, we show that the recombination event between *Satellite* and *Independent* alleles is much more recent than previously thought and likely occurred within the past 100,000 years. Our comprehensive description of the evolution of the ruff mating polymorphism demonstrates how supergenes evolve over time and how they contribute to the maintenance of genetic variation and phenotypic evolution in nature.

## Author contributions

L.A. conceived the study. F.W. and S.L. performed field work and collected tissues samples. D.S. prepared genomic DNA and mRNA. S.Y. prepared the RNAseq library, H.B. contributed carried out experimental work. J.H. and E.D.E. were responsible for the bioinformatic analysis. J.H., E.D.E., and L.A. wrote the paper with input from other authors. All authors approved the manuscript before submission.

## Data availability statement

The data have been submitted to the short reads archive (http://www.ncbi.nlm.nih.gov/sra) under BioProject PRJNA816664.

## Code availability statement

The analyses of data have been carried out with publicly available software and all are cited in the Methods section. Code associated with bioinformatic analyses are available at: https://github.com/LeifAnderssonLab/Ruff_assembly_2022.

## Acknowledgements

We thank Russel Corbett-Detig, Peter Grant and Rosemary Grant for comments on the manuscript and Mats Pettersson for advice concerning the HaploDistScan analysis. The project was financially supported by Vetenskapsrådet (2017-02907) and Knut and Alice Wallenberg Foundation (KAW 2016.0361). The National Genomics Infrastructure (NGI)/Uppsala Genome Center provided service in massive parallel sequencing and the computational infrastructure was provided by the Swedish National Infrastructure for Computing (SNIC) at UPPMAX partially funded by the Swedish Research Council (2018-05973).

## Author information

The authors declare no competing financial interests. Correspondence and requests for materials should be addressed to L.A.

## Supplementary Figures

**Supplementary Fig. 1.**
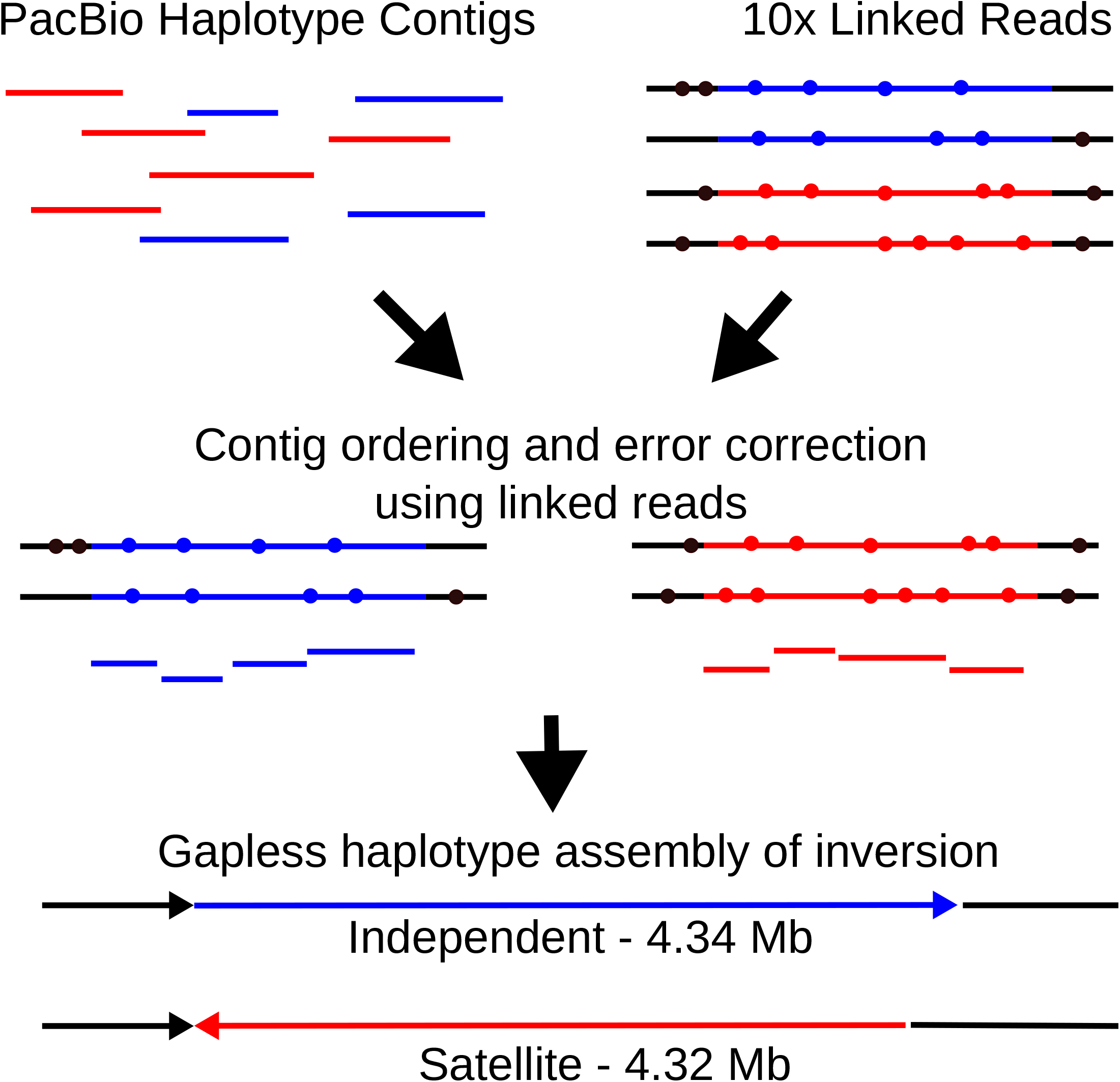
Inversion assembly methods. Satellite (inverted) and Independent (non-inverted) haplotypes were separately assembled using contigs from a PacBio Falcon v0.5 diploid assembly. These contigs were phased and ordered by alignment to a Supernova v2.0 whole genome diploid assembly based on Chromium 10x linked reads. The assembled haplotypes were then polished with Chromium 10x linked read sequences from the same haplotype. The final Satellite assembly contained no additional large structural differences, however the accumulation of small deletions in the Satellite haplotype resulted in an inversion length 14 kb shorter than the corresponding Independent sequence.

**Supplementary Fig. 2.**
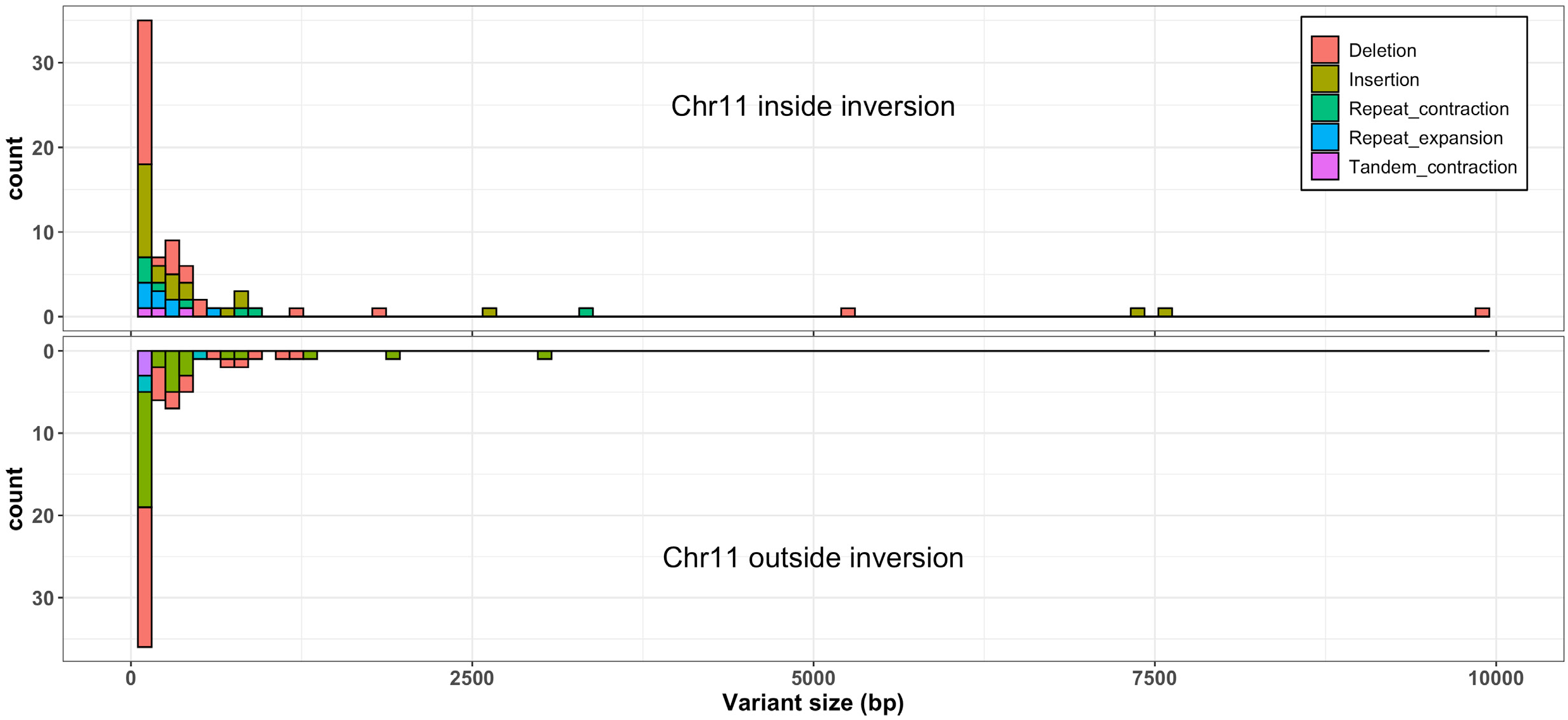
The Satellite inversion haplotype is not enriched for repeats. Above, the size distribution of deletions, insertions, and repetitive elements in the Satellite inversion haplotype. Below, the size distribution of deletions, insertions, and repetitive elements in the remainder of chromosome 11. Contractions and expansions are annotated relative to the Independent chromosome. The Satellite allele does not show an enrichment of repetitive elements neither compared to the Independent haplotype nor relative to the remainder of chromosome 11.

**Supplementary Fig. 3.**
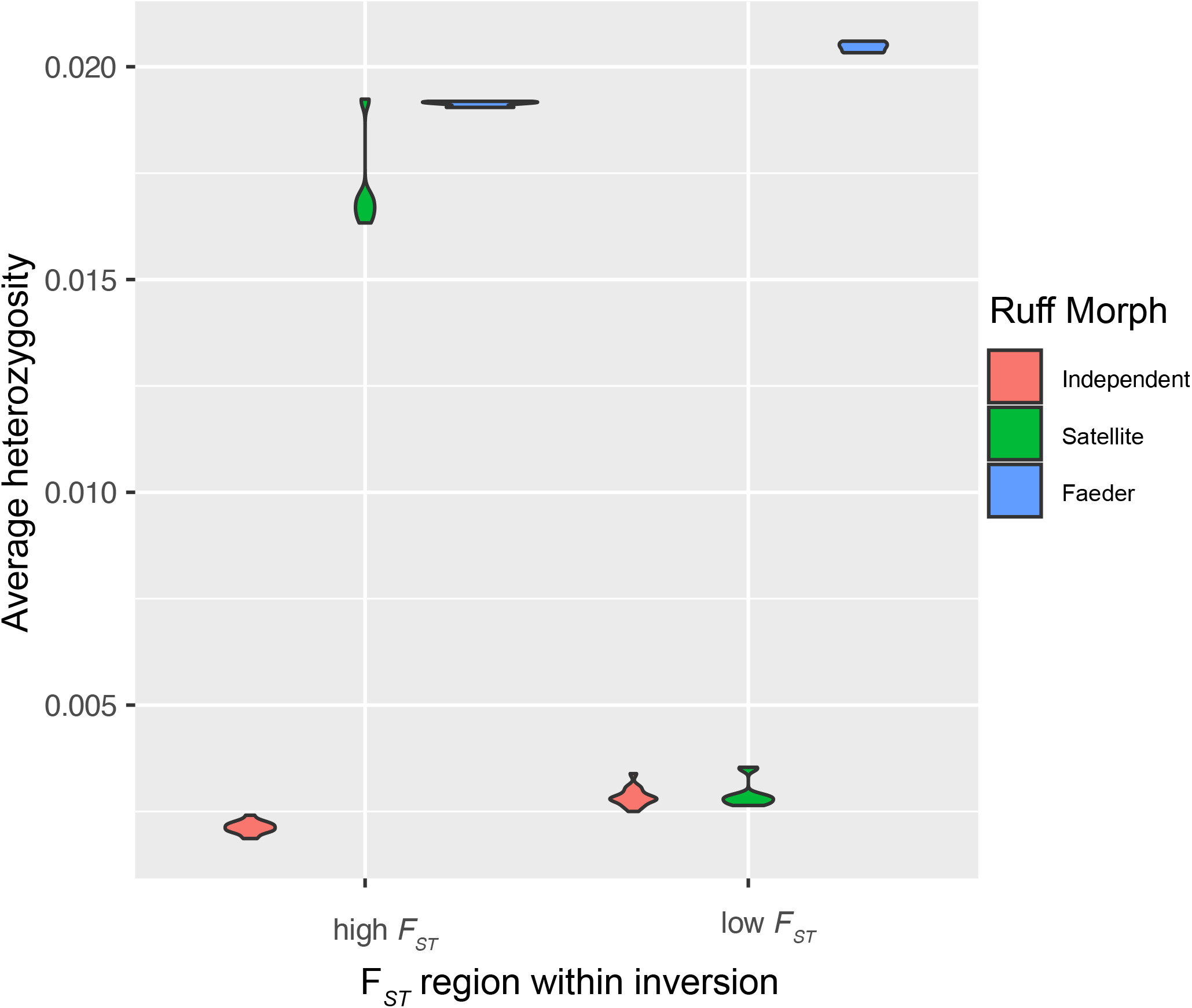
Nucleotide diversity in Satellites and Independents. The average heterozygosity within the high (red) and low (blue) *F_ST_* regions was calculated for each resequenced ruff individual. The only Faeder individual show high levels of heterozygosity throughout the inversion. Heterozygosity levels between Independent and Satellite samples are indistinguishable in the low *F_ST_* regions indicating a recent haplotype divergence. Based on these data we estimated the pairwise nucleotide diversity (*dxy*) between Satellite and Independent haplotypes at 0.0170 and 0.0028 for the high and low *F_ST_* regions, respectively; and the average nucleotide diversity among Independent haplotypes at 0.0022 and 0.0029 for the high and low *F_ST_* regions, respectively.

**Supplementary Fig. 4.**
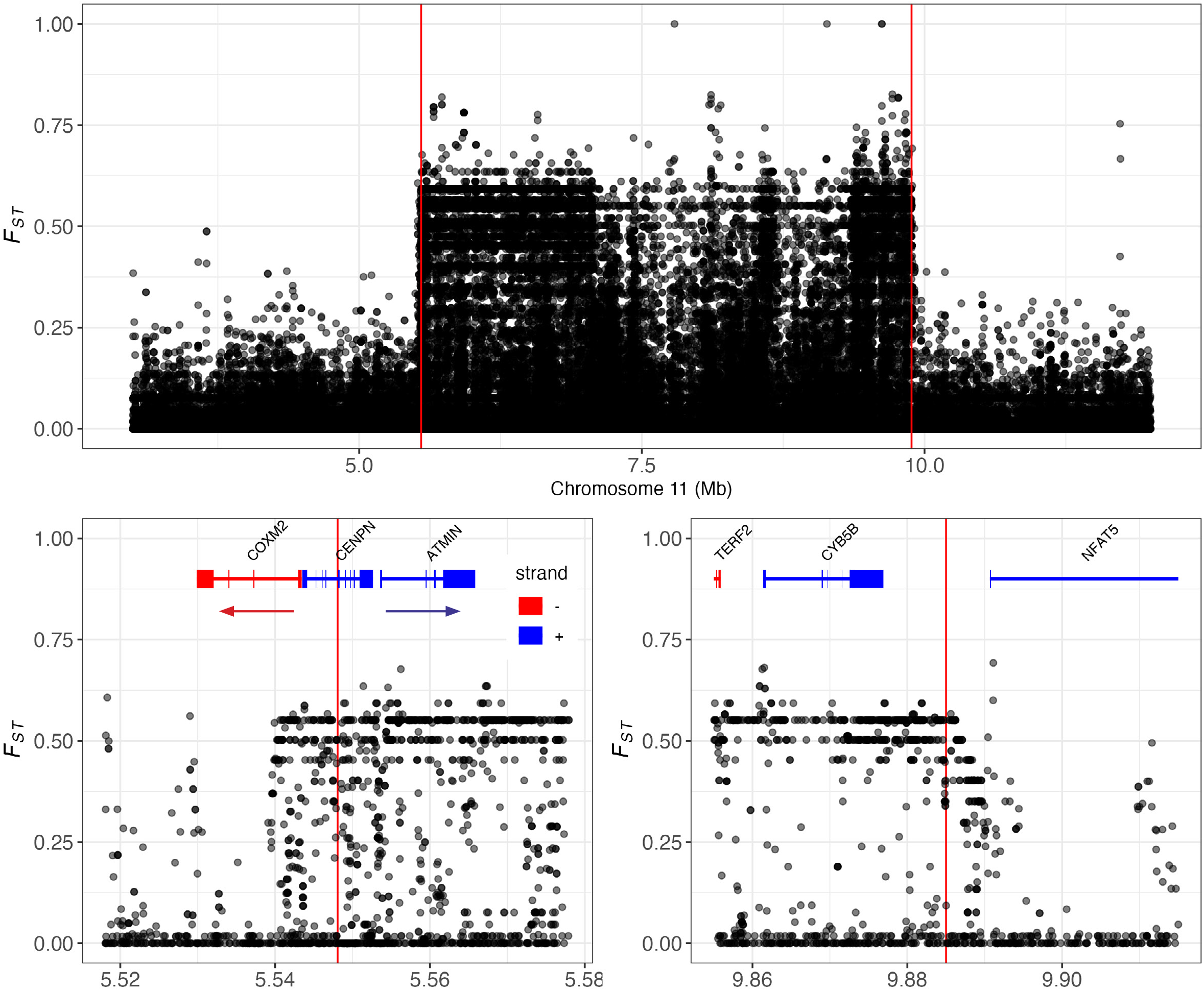
Per-site genetic divergence across chromosome 11. Above, per-site *F_ST_* between Satellite and Independent individuals across the genomic region of chromosome 11 harboring the inversion. Below, per-site *F_ST_* for the 5.55 Mb and 9.89 Mb breakpoints. Note the disruption of *CENPN* at the 5.55 Mb breakpoint. Elevated *F_ST_* upstream and downstream of the inverted region suggests linkage disequilibrium extends beyond the inverted region (see also Fig. 3).

**Supplementary Fig. 5.**
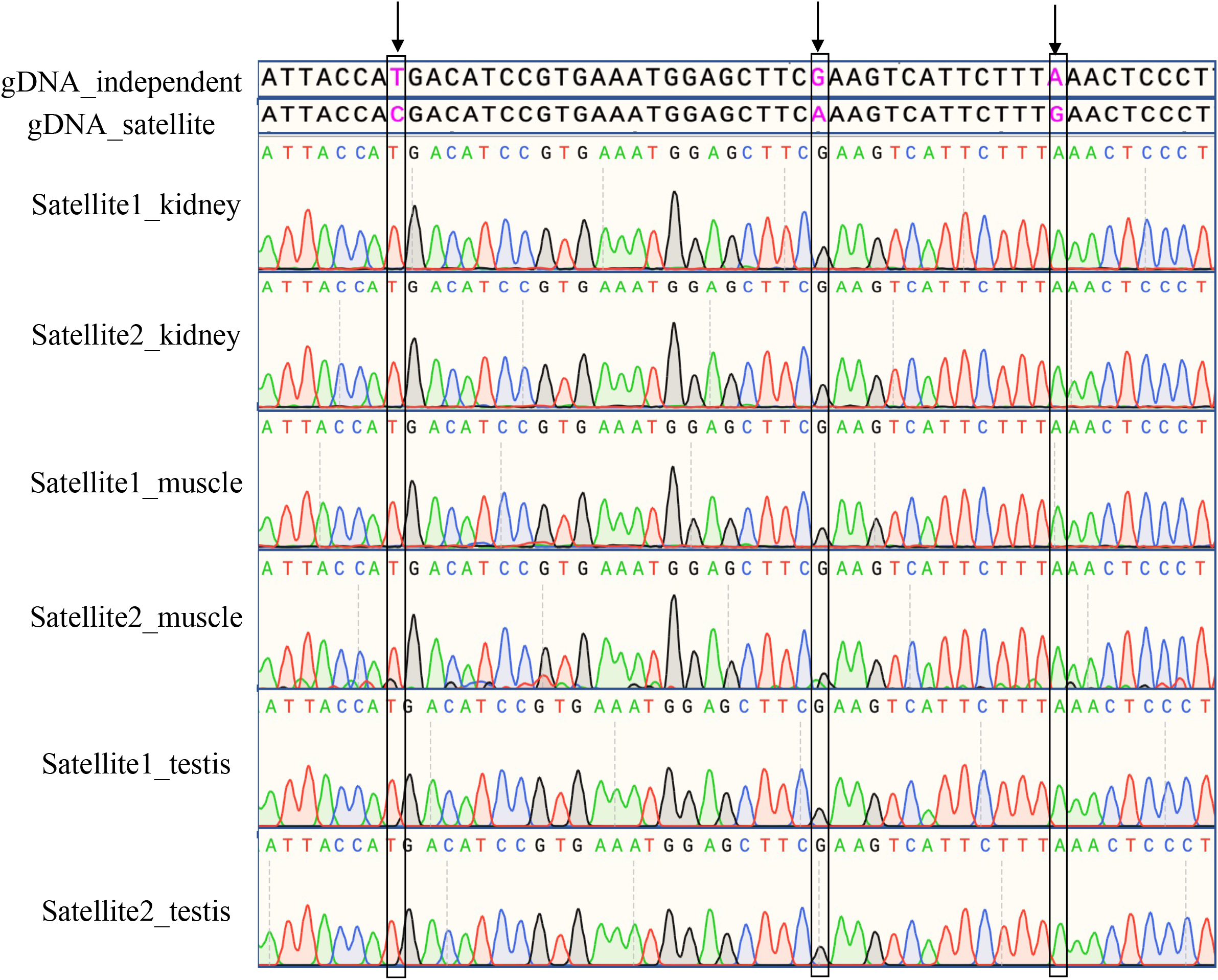
RT-PCR analysis of *CENPN* expression. Genomic DNA sequence from Independent and Satellite are on top, corresponding cDNA sequence for the parts of the *CENPN* transcript encoded by exons inside the inversion are on the bottom. Three informative SNPs are highlighted by black arrow and box.

**Supplementary Fig. 6.**
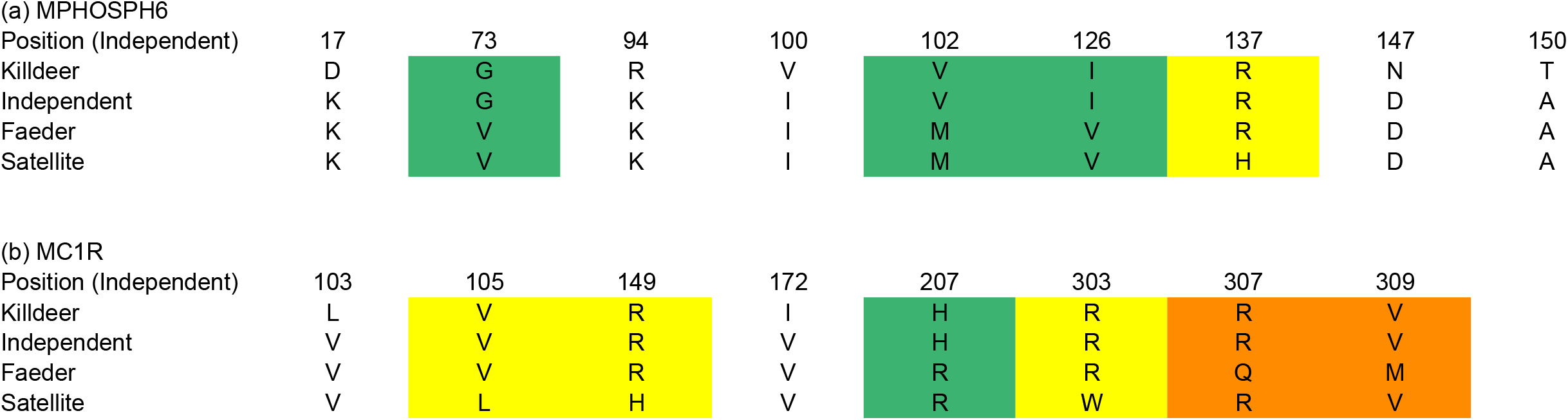
Amino acid alignment of variable sites in MC1R (**a**) and MPHOSP6 (**b**) among killdeer and the three different ruff morphs.

**Supplementary Fig. 7.**
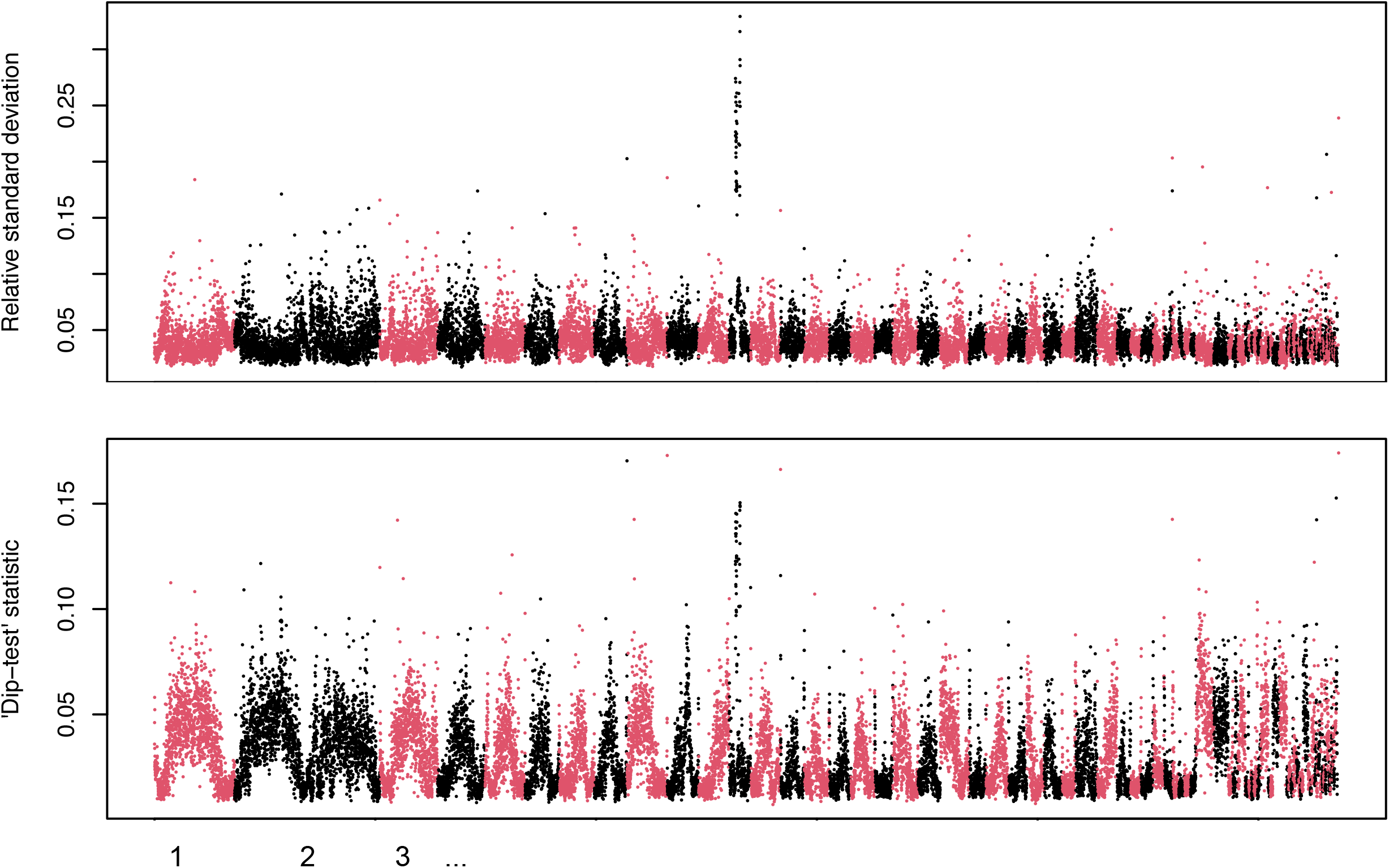
Sequence divergence within all ruff individuals across the genome based on the HaploDistScan method^35^. This method searches the genome for the presence of divergent haplotypes, such as the segregating haplotypes in the ruff supergene. The two plots show two summary statistics used to describe haplotype divergence within 10 kb windows across the genome. a) contains the standard deviation of genetic distance among all haplotypes in the sample and b) contains the ‘dip test’ statistics based on the modality of sequence divergence within the target region. The chr11 inversion is the only region with consistent elevated divergence among haplotypes. Several individual windows are elevated in the dip test statistic, but are not strongly elevated in panel a) containing estimated genetic distances among haplotypes in the region. This suggests that many of the individual elevated windows detected by the dip test are not segregating for haplotypes as old as the inversion and are not likely candidates for being remnant introgressed fragments from another species. The result is not unexpected even if the inversion originated due to an interspecies introgression event, because neutral alleles in other parts of the genome will be lost after many generations of back-crossing to ruff. This may occur at an accelerated rate as the inversion is a recessive lethal.

## Supplementary Tables

**Supplementary Table 1.** Genome assembly statistics generated by Supernova using Chromium 10x linked reads.

|  |  |
| --- | --- |
| Assembly size (bp) | 1,129,272,262 |
| Mean bases per read | 138.5 |
| N50 contig length | 294,031 |
| Percent duplicate | 22.79 |
| N50 edge length | 30,656 |
| Effective read coverage | 77.77 |
| Effective read coverage, median definition | 79.23 |
| Estimated genome size (bp) | 1,296,440,000 |
| Mean distance between heterozygous SNPs | 358 |
| Length-weighted mean molecule length | 43,717.6 |
| Percent of base assembly missing from final assembly in scaffolds $\geq 10$ kb | 4.84 |
| Median insert size | 322 |
| Number of reads | 1,050,579,400 |
| Average number of molecules extending 10 kb in both directions from a given point | 157.5 |
| N50 phase block length | 20,185,970 |
| Nonduplicate and phased reads | 48.06 |
| Percent proper pairs | 89.78 |
| Raw coverage | 121.55 |
| Repetitive fraction | 5.56 |
| N50 reads per barcode | 866 |
| N50 scaffold length | 27,708,464 |
| Scaffolds $\geq 10$ kb | 697 |
| Scaffolds $\geq 1$ kb | 10,826 |
| Percent of reads that are not barcoded | 5.48 |
| Percent reads with valid barcodes | 94.52 |

Genome assembly statistics for Ruff Satellite individual using 10x linked reads with the Supernova assembler v.2.0. Additional work was done to manually assemble the inversion using PacBio long reads, however the overall genome was assembled according to the above specifications.

**Supplementary Table 2.** Current gene annotation summary of the inversion region and comparison to previous annotation of scaffold 28. (Excel file).

**Supplementary Table 3.** Summary of dNdS statistics of the inversion genes compared between killdeer and ruff morphs. Excel file.

## Methods

### Genome assembly

A male ruff individual with the Satellite phenotype was collected in northern Sweden (coordinates WGS 84 (latitude, longitude) 68.1, 19.8) during the reproductive season of 2016 under permit Swedish Environmental Protection Agency NV 02900-16. Small muscle pieces were stored in RNAlater (ThermoFisher) at 4°C until DNA preparation using the DNeasy Blood and Tissue Kit (Qiagen) according to the manufacture’s recommendation. A Chromium 10x linked read library was produced according to the manufacturer’s recommendation and sequenced on an Illumina HiSeqX to a target depth of 90x. Supernova (v2.0)^36^ was used to build a diploid assembly with a phased contig N50 of 20.2 Mb. A PacBio long read library was prepared from the same DNA sample according to the manufacture’s recommendation and processed with Falcon (v0.5)^37^ to produce a second diploid genome assembly. Assembly scaffolds were aligned to *G. gallus* and *C. pugnax* genomes using Satsuma2 (v2016-12-07, https://github.com/bioinfologics/satsuma2) chromosemble to identify the scaffold homologous to the part of chicken chromosome 11 which correspond to the region of the ruff genome harboring the inversion. Chromium 10x linked reads were mapped to the genome assembly using longranger (v2.2.2)^36^ and PacBio long reads were mapped with minimap2 (v2.14)^38^. Longranger identified structural variants which together with manual inspection using IGV (v 2.5.3)^39^ revealed the inversion breakpoints.

While the Supernova assembly using linked reads provided a contiguous and well phased genome assembly the nature of the short-read genome assembly process left inevitable gaps in the sequence. To produce a complete assembly of both the Independent and Satellite haplotype of the inversion region the Falcon long read assembly contigs were identified as Independent or Satellite using the phased SNPs from the supernova assembly. Falcon contigs were then aligned to each-other using Satsuma2 and BLAST (v2.11.0)^40^ to resolve overlaps. Incorrect haplotype switching within Falcon contigs was identified using Chromium 10x linked reads and manually corrected. Corrected Falcon contigs were merged into gapless haplotype assemblies and polished using Pilon (v1.22)^41^ and the Chromium 10x reads split by haplotype.

### Comparison to previous short-read assembly

Chromosome 11 of the genome assembly reported above was aligned to Scaffold 28 of the previous short-read assembly^3^ using the MUMmer (v4.0.0rc1)^42^ dnadiff tool with default parameters. Gaps identified between the present assembly and the former short-read assembly in the inversion region were intersected with between-contig stretches of Ns in the short-read assembly using BEDTools (2.29.2)^43^. This exercise provided an estimate of the assembly size inflation due to the addition of Ns during scaffolding of the short-read assembly. Long PacBio reads mapped to the current and previous assemblies as described above were used to determine support for apparent sequence duplication in the short read assembly as well as for visual inspection of uncategorized discrepancies greater than 1 kb.

### Genome alignments

Satellite and Independent chromosome 11 were aligned to each other using MUMmer (nucmer)^42^. The corresponding alignments were uploaded to the Assemblytics^44^ web portal and variants called with unique sequence length = 10,000, maximum variant size = 10,000, and minimum variant size = 50.

### Genome annotation

We annotated the Independent and Satellite haplotypes (of chromosome 11) using MAKER (3.01.2-beta)^45^. Prior to annotation, we created a custom repeat library using Repeat Modeler (1.0.8, http://www.repeatmasker.org) and RepeatMasker (4.0.7, http://www.repeatmasker.org) and downloaded all protein sequences in the curated and reviewed Swiss-Prot database^46^ and for the previous assembly of Ruff (GCF_001431845). We downloaded RNAseq reads for 10 previously sequenced individuals^18^ across 5 tissues and additionally included a single skin tissue from the present study. Briefly, skin tissue was dissected in the field and immediately stored in RNAlater (ThermoFisher) to stabilize the RNA. RNA was extracted and DNase treated as described previously^47^. The RNA quality and concentration were measured by the RNA ScreenTape assay (TapeStation, Agilent Technologies). Strand-specific mRNA sequencing libraries were generated using the SENSE RNA-Seq Library Prep kit (Lexogen). Briefly, 1 μg of total RNA was poly-A selected using magnetic beads. Illumina-compatible linker sequences were introduced to the mRNA by random hybridization. The amplified libraries were size-selected for an average insert size of ~350 bp and sequenced using an Illumina HiSeq instrument at SciLifeLab-Uppsala, Sweden.

We mapped RNAseq reads to each genome assembly containing either the Independent or Satellite chromosome using HISAT2^48^ and assembled transcripts using StringTie^49^. We extracted splice junctions by mapping all reads with TopHat2^50^ and converting to gff3 with MAKER’s^45^ tophat2gff3 script. To generate high quality gene models on each chromosome version, we ran MAKER using protein and RNAseq data as evidence, splice junction annotations, and lists of repeats to mask low confidence regions. We ran MAKER with best practices recommendations and additionally included max_dna_len=150000 and split_hit=50000 to fine tune annotation for an avian genome. All MAKER-predicted proteins were blasted (using BLAST 2.7.1+)^40^ against the given protein evidence to produce candidate ortholog annotations for each annotated gene and the resulting gff updated using maker_functional_gff.

Manual curation of the gene models was carried out using the web-based genomic annotation editing platform Web Apollo^51^ for both the Independent and Satellite allele of each gene. An exon was included in a gene isoform if it was supported by at least three RNA-seq reads with identical splice boundaries in an individual. Exon boundaries were defined by the longest continuous block of RNA-seq reads. The longest isoform was chosen as the representative of a gene in further analysis.

### Nucleotide substitution rates

The manually curated gene set was used to calculate nucleotide substitution rates between Independent and Satellite alleles of each gene and their orthologs in killdeer. The killdeer gene set (Genbank ID: 1184028) was downloaded and 45 orthologs present within the high *F_ST_* regions in the comparison of Satellite and Independent haplotypes were identified using reciprocal best blast between protein sequences. All ortholog and allele pairs were aligned using ClustalO (v1.2.4)^52^ and alignments given to PAML (v4.9e)^23^ CODEML to calculate dN and dS values for each pair.

Relaxed purifying selection is expected for many genes on the Satellite haplotype because this haplotype always occurs in the presence of Independent haplotypes in *S/I* heterozygotes. The deviation of the observed dN on the Satellite haplotype from expected dN in the absence of purifying selection was calculated by estimating the increase in expected nonsynonymous substitution on the Satellite haplotype, ΔdNs, as follows.

Definitions

dNs = dN in the Satellite branch since formation of the inversion
dNi = dN in the Independent branch since formation of the inversion
dNis = dN between Independent and Satellite
dSis = dS between Independent and Satellite

The amount dN increases in Satellites if there is no purifying selection can be estimated by

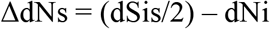

However, since we do not know the value of dNi directly we can estimate dNi based on dNis and dSis:

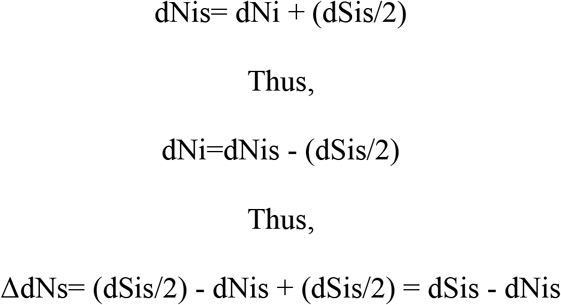

Based on this we calculated expected dN (between Satellite ruff vs. killdeer) as dN (between Independent ruff vs. killdeer) + ΔdNs and compared this with observed dN (between Satellite ruff vs. killdeer).

The CODEML function of the PAML program^23^ was used to test the relative likelihood of models in which substitution rates varied on the branches leading to the ruff orthologs of *CENPN, MC1R*, and *MPHOSPH6*. For each gene the orthologs for ruff, killdeer, golden eagle, chicken, and mallard were aligned as described above in the section on dN/dS. CODEML then calculated the log-likelihood for models in which the substation rate ω (*dN/dS*) was held constant for the entire ortholog tree (model 0), or allowed to vary only on the branch leading to Satellite ruff haplotype (model S). The significance of the difference between the log-likelihoods of the two models was calculated with the Pearson chi-squared test using a one tailed distribution.

### Genetic variation analysis

Individual resequencing data from ruff were downloaded from two previous studies: 25 individuals from Lamichhaney *et al*.^3^, and 5 individuals from Küpper *et al.*^18^. Read sets were adapter trimmed with bbmap v38.61b (ktrim=r, qout=33, k=23, mink=11, hdist=1, qtrim=r, trimq=10, maq=10, tpe, ow=t, tbo) and mapped to the Independent ruff genome assembly using BWA-MEM (v0.7.17)^53^ and default settings. Biallelic SNPs were called using GATK (v4.1.1.0)^54^ following GATK best practices for non-model organisms (see code availability). VCF files were analyzed in R (v4.0.3)^55^ using the packages zoo (v1.8-8)^56^, viridis (v0.6.1)^57^, tidyverse (v1.3.1)^58^, and vcfr (v1.12.0)^59^ within custom scripts (Supplementary Data 1, (see code availability).

### *F_ST_* calculations

We estimated relative divergence (*F_ST_*) using the software *pixy*^60^ in 15-kb non-overlapping sliding windows and per site by setting the window size in *pixy* to 1-bp.

### Estimating the age of the inversion and the Satellite haplotype

Nucleotide diversity among Satellite chromosomes cannot be calculated using standard software because these only occur in the heterozygous state. We therefore estimated *d_XY_* between Satellite and Independent chromosomes based on the average number of heterozygous sites among callable sites in Satellite males (*S/I*) and nucleotide diversity (*d_X_*) among Independents (*I/I*) in the same fashion. The classical way to calculate divergence time between DNA sequences from two populations is based on calculating the net number of nucleotide substitutions taking account the intrapopulation nucleotide diversities using the formula *d_a_* = *d_XY_* – (*d_X_* + *d_Y_*)/2 according to Nei^61^; *d_X_* and *d_Y_* are the intrapopulation nucleotide differences within populations X and Y. However, this is not applicable here because the Satellite and Independent chromosomes originates from the same population and therefore a correct estimate is obtained using *d_a_* = *d_XY_* - *d_X_*; *d_X_* are the intrapopulation nucleotide differences among Independent chromosomes.

Time since divergence between Satellite and Independent chromosomes was estimated using T= *d_a_*/2λ and among Independent chromosomes as T=*d_x_*/2 λ according to Nei^61^ where λ is the average nucleotide substitution rate per year and here we used the average estimate for birds 1.9 x 10-^9^ (Ref.^62^).

### Introgression analysis

Biallelic SNPs generated above were further filtered to exclude any sites with missing samples. This filtered SNP set was phased for the whole genome using BEAGLE (5.4)^63^ with default parameters. Phased genotypes were then analysed using HaploDistScan^35^.

### *CENPN* RT-PCR analysis

Total RNA was isolated from testis, kidney and muscle of two Satellite and two Independent individuals as described above for RNA-seq. First-strand cDNA was synthesized using High-Capacity cDNA Reverse Transcription Kit (Thermo Scientific). The parts of the *CENPN* transcript encoded by exons inside the inversion were amplified using the following primers (5’-3’): F: AGGATGTGGTTTATCTTTGTGAGGAAA; R: TCTCAAGCCTATATTGTGCAAATTC. The amplification program was as follows: 95°C for 5min, followed by 35 cycles of 95°C for 30s, 60°C for 30s and 72°C for 1min. PCR products were Sanger sequenced.

## Literature

1 Widemo, F. Alternative reproductive strategies in the ruff: a mixed ESS? Anim. Behav. 13, 329–336 (1998).

2 Jukema, J. & Piersma, T. Permanent female mimics in a lekking shorebird. Biol. Lett. 2, 161–164 (2006).

3 Lamichhaney, S., Fan, G., Widemo, F., Gunnarsson, U. & Schwochow Thalmann, D. Structural genomic changes underlie alternative reproductive strategies in the ruff *(Philomachus pugnax)* Nat Gen 48, 84–88 (2016).

4 Gutiérrez-Valencia, J., Hughes, P. W., Berdan, E. L. & Slotte, T. The genomic architecture and evolutionary fates of supergenes. Genome Biol Evol 13, evab057 (2021).

5 Faria, R., Johannesson, K., Butlin, R. K. & Westram, A. M. Evolving inversions. Trends Ecol Evol 34, 239–248, doi:10.1016/j.tree.2018.12.005 (2019).

6 Linksvayer, T. A., Busch, J. W. & Smith, C. R. Social supergenes of superorganisms: do supergenes play important roles in social evolution? Bioessays 35, 683–689, doi:10.1002/bies.201300038 (2013).

7 Kirkpatrick, M. & Barton, N. Chromosome inversions, local adaptation and speciation. Genetics 173, 419–434, doi:10.1534/genetics.105.047985 (2006).

8 Nishikawa, H. et al. A genetic mechanism for female-limited Batesian mimicry in Papilio butterfly. Nat Genet 47, 405–409, doi:10.1038/ng.3241 http://www.nature.com/ng/journal/v47/n4/abs/ng.3241.html#supplementary-information (2015).

9 Brelsford, A. et al. An ancient and eroded social supergene is widespread across Formica ants. Curr Biol 30, 304–311.e304, doi:10.1016/j.cub.2019.11.032 (2020).

10 Pearse, D. E. et al. Sex-dependent dominance maintains migration supergene in rainbow trout. Nat Ecol Evol 3, 1731–1742, doi:10.1038/s41559-019-1044-6 (2019).

11 Thomas, J. W. et al. The chromosomal polymorphism linked to variation in social behavior in the white-throated sparrow *(Zonotrichia albicollis)* is a complex rearrangement and suppressor of recombination. Genetics 179, 1455–1468, doi:10.1534/genetics.108.088229 (2008).

12 Han, F. et al. Ecological adaptation in Atlantic herring is associated with large shifts in allele frequencies at hundreds of loci. eLife 9, e61076, doi:10.7554/eLife.61076 (2020).

13 Matschiner, M. et al. Supergene origin and maintenance in Atlantic cod. Nat Ecol Evol, doi: 10.1038/s41559-022-01661-x (2022).

14 Charlesworth, B. & Charlesworth, D. Evolution: a new idea about the degeneration of Y and W chromosomes. Curr Biol 30, R871–r873, doi:10.1016/j.cub.2020.06.008 (2020).

15 Stolle, E. et al. Degenerative expansion of a young supergene. Mol Biol Evol 36, 553–561, doi:10.1093/molbev/msy236 (2019).

16 Berdan, E. L., Blanckaert, A., Butlin, R. K. & Bank, C. Deleterious mutation accumulation and the long-term fate of chromosomal inversions. PLoS Genet 17, e1009411, doi:10.1371/journal.pgen.1009411 (2021).

17 Navarro, A., Betrán, E., Barbadilla, A. & Ruiz, A. Recombination and gene flux caused by gene conversion and crossing over in inversion heterokaryotypes. Genetics 146, 695–709, doi:10.1093/genetics/146.2.695 (1997).

18 Küpper, C. et al. A supergene determines highly divergent male reproductive morphs in the ruff. Nat. Genet. 48, 79–83 (2016).

19 Lank, D. B., Smith, C. M., Hanotte, O., Burke, T. & Cooke, F. Genetic polymorphism for alternative mating behaviour in lekking male ruff Philomachus pugnax. Nature 378, 59–62 (1995).

20 Lank, D. B., Farrell, L. L., Burke, T., Piersma, T. & McRae, S. B. A dominant allele controls development into female mimic male and diminutive female ruffs. Biol. Lett. 9, 20130653 (2013).

21 Höglund, J. & Lundberg, A. Plumage color correlates with body size in the ruff (*Philomachus pugnax*). Auk 13, 306–308 (1989).

22 Jay, P. et al. Mutation load at a mimicry supergene sheds new light on the evolution of inversion polymorphisms. Nat Genet 53, 288–293, doi:10.1038/s41588-020-00771-1 (2021).

23 Yang, Z. PAML 4: a program package for phylogenetic analysis by maximum likelihood. Mol. Biol. Evol. 24, 1586–1591 (2007).

24 Loveland, J. L., Lank, D. B. & Küpper, C. Gene expression modification by an autosomal inversion associated with three male mating morphs. Front Genet 12, 641620, doi:10.3389/fgene.2021.641620 (2021).

25 Prum, R. O. et al. A comprehensive phylogeny of birds (Aves) using targeted nextgeneration DNA sequencing. Nature 526, 569–573, doi:10.1038/nature15697 (2015).

26 Clarke, C. A. & Sheppard, P. M. Super-genes and mimicry. Heredity 14, 175–185, doi:10.1038/hdy.1960.15 (1960).

27 Matsumoto-Taniura, N., Pirollet, F., Monroe, R., Gerace, L. & Westendorf, J. M. Identification of novel M phase phosphoproteins by expression cloning. Mol Biol Cell 7, 1455–1469, doi:10.1091/mbc.7.9.1455 (1996).

28 Schilders, G., Raijmakers, R., Raats, J. M. & Pruijn, G. J. MPP6 is an exosome-associated RNA-binding protein involved in 5.8S rRNA maturation. Nucleic Acids Res 33, 6795–6804, doi:10.1093/nar/gki982 (2005).

29 Li, C. et al. Genome-wide association analysis in humans links nucleotide metabolism to leukocyte telomere length. Am J Hum Genet 106, 389–404, doi:10.1016/j.ajhg.2020.02.006 (2020).

30 Imsland, F. et al. The Rose-comb mutation in chickens constitutes a structural rearrangement causing both altered comb morphology and defective sperm motility. PLoS Genet. 8, e1002775, doi:Artn E1002775 Doi 10.1371/Journal.Pgen.1002775 (2012).

31 Schwander, T., Libbrecht, R. & Keller, L. Supergenes and complex phenotypes. Curr Biol 24 (2014).

32 Gibson, R. & Baker, A. Multiple gene sequences resolve phylogenetic relationships in the shorebird suborder Scolopaci (Aves: Charadriiformes). Molecular Phylogenetics and Evolution 64, 66–72, doi:https://doi.org/10.1016/j.ympev.2012.03.008 (2012).

33 Corbett-Detig, R. B. & Hartl, D. L. Population genomics of inversion polymorphisms in *Drosophila melanogaster*. PLOS Genetics 8, e1003056, doi:10.1371/journal.pgen.1003056 (2012).

34 Marklund, L., Moller, M. J., Sandberg, K. & Andersson, L. A missense mutation in the gene for melanocyte-stimulating hormone receptor (*MC1R*) is associated with the chestnut coat color in horses. Mamm. Genome 7, 895–899 (1996).

35 Pettersson, M. E., Kierczak, M., Almén, M. S., Lamichhaney, S. & Andersson, L. A model-free approach for detecting genomic regions of deep divergence using the distribution of haplotype distances. bioRxiv, 144394, doi:10.1101/144394 (2017).

36 Weisenfeld, N. I., Kumar, V., Shah, P., Church, D. M. & Jaffe, D. B. Direct determination of diploid genome sequences. Genome Res 27, 757–767, doi:10.1101/gr.214874.116 (2017).

37 Chin, C. S. et al. Phased diploid genome assembly with single-molecule real-time sequencing. Nat Methods 13, 1050–1054, doi:10.1038/nmeth.4035 (2016).

38 Li, H. Minimap2: pairwise alignment for nucleotide sequences. Bioinformatics 34, 3094–3100, doi:10.1093/bioinformatics/bty191 (2018).

39 Robinson, J. T. et al. Integrative Genomics Viewer. Nat. Biotech. 29, 24–26 (2011).

40 Camacho, C. et al. BLAST+: architecture and applications. BMC Bioinformatics 10, 421, doi:10.1186/1471-2105-10-421 (2009).

41 Walker, B. J. et al. Pilon: an integrated tool for comprehensive microbial variant detection and genome assembly improvement. PLOS ONE 9, e112963, doi:10.1371/journal.pone.0112963 (2014).

42 Marçais, G. et al. MUMmer4: A fast and versatile genome alignment system. PLoS Comput Biol 14, e1005944, doi:10.1371/journal.pcbi.1005944 (2018).

43 Quinlan, A. R. & Hall, I. M. BEDTools: a flexible suite of utilities for comparing genomic features. Bioinformatics 26, 841–842, doi:10.1093/bioinformatics/btq033 (2010).

44 Nattestad, M. & Schatz, M. C. Assemblytics: a web analytics tool for the detection of variants from an assembly. Bioinformatics 32, 3021–3023, doi:10.1093/bioinformatics/btw369 (2016).

45 Cantarel, B. L. et al. MAKER: An easy-to-use annotation pipeline designed for emerging model organism genomes. Genome Res. 18, 188–196, doi:10.1101/gr.6743907 (2008).

46 Consortium, T. U. UniProt: the universal protein knowledgebase in 2021. Nucleic Acids Research 49, D480–D489, doi:10.1093/nar/gkaa1100 (2020).

47 Schwochow Thalmann, D. et al. The evolution of *Sex-linked barring* alleles in chickens involves both regulatory and coding changes in *CDKN2A*. PLoS Genet 13, e1006665 (2017).

48 Kim, D., Paggi, J. M., Park, C., Bennett, C. & Salzberg, S. L. Graph-based genome alignment and genotyping with HISAT2 and HISAT-genotype. Nat Biotechnol 37, 907–915, doi:10.1038/s41587-019-0201-4 (2019).

49 Pertea, M., Kim, D., Pertea, G. M., Leek, J. T. & Salzberg, S. L. Transcript-level expression analysis of RNA-seq experiments with HISAT, StringTie and Ballgown. Nat Protoc 11, 1650–1667, doi:10.1038/nprot.2016.095 (2016).

50 Kim, D. et al. TopHat2: accurate alignment of transcriptomes in the presence of insertions, deletions and gene fusions. Genome Biol. 14, R36 (2013).

51 Lee, E. et al. Web Apollo: a web-based genomic annotation editing platform. Genome Biology 14, R93, doi:10.1186/gb-2013-14-8-r93 (2013).

52 Sievers, F. et al. Fast, scalable generation of high-quality protein multiple sequence alignments using Clustal Omega. Mol Syst Biol 7, 539, doi:10.1038/msb.2011.75 (2011).

53 Li, H. Aligning sequence reads, clone sequences and assembly contigs with BWA-MEM. arXiv:1303.3997v2 [q-bio.GN] (2013).

54 Poplin, R. et al. Scaling accurate genetic variant discovery to tens of thousands of samples. bioRxiv, 201178, doi:10.1101/201178 (2018).

55 Team, R. C. R: A language and environment for statistical computing. R Foundation for Statistical Computing, Vienna, Austria. (2020).

56 Zeileis, A. & Grothendieck, G. zoo: S3 Infrastructure for Regular and Irregular Time Series. Journal of Statistical Software 14, 1–27, doi:10.18637/jss.v014.i06 (2005).

57 Garnier, S. et al. (2021). viridis - Colorblind-Friendly Color Maps for R. doi: 10.5281/zenodo.4679424, R package version 0.6.2, https://sjmgarnier.github.io/viridis/ (2021).

58 Wickham, H. et al. Welcome to the Tidyverse Journal of Open Source Software 4, 1686 (2019).

59 Knaus, B. J. & Grünwald, N. J. vcfr: a package to manipulate and visualize variant call format data in R. Mol Ecol Resour 17, 44–53, doi:10.1111/1755-0998.12549 (2017).

60 Korunes, K. L. & Samuk, K. pixy: Unbiased estimation of nucleotide diversity and divergence in the presence of missing data. Molecular Ecology Resources 21, 1359–1368, doi:https://doi.org/10.1111/1755-0998.13326 (2021).

61 Nei, M. Molecular Evolutionary Genetics. 276–279 (Columbia University Press, 1987).

62 Zhang, G. et al. Comparative genomics reveals insights into avian genome evolution and adaptation. Science 346, 1311–1320 (2014).

63 Browning, B. L., Tian, X., Zhou, Y. & Browning, S. R. Fast two-stage phasing of large-scale sequence data. The American Journal of Human Genetics 108, 1880–1890, doi:https://doi.org/10.1016/j.ajhg.2021.08.005 (2021).

